# Evaluation of the nemabiome approach for the study of equine strongylid communities

**DOI:** 10.1101/2022.07.22.501098

**Authors:** Élise Courtot, Michel Boisseau, Sophie Dhorne-Pollet, Delphine Serreau, Amandine Gesbert, Fabrice Reigner, Marta Basiaga, Tetiana Kuzmina, Jérôme Lluch, Gwenolah Annonay, Claire Kuchly, Irina Diekmann, Jürgen Krücken, Georg von Samson-Himmelstjerna, Nuria Mach, Guillaume Sallé

## Abstract

Basic knowledge on the biology and epidemiology of equine strongylid species remains insufficient although it would contribute to the design of better parasite control strategies. Nemabiome is a convenient tool to quantify and to identify species in bulk samples that could overcome the hurdle that cyathostomin morphological identification represents. To date, this approach has relied on the internal transcribed spacer 2 (ITS-2) of the ribosomal RNA cistron and its predictive performance and associated biases both remain unaddressed.

This study aimed to bridge this knowledge gap using cyathostomin mock communities and comparing performances of the ITS-2 and a *cytochrome c oxidase subunit I* (COI) barcode newly developed in this study. The effects of bioinformatic parameters were investigated to determine the best analytical pipelines. Subsequently, barcode predictive abilities were compared across various mock community compositions. The replicability of the approach and the amplification biases of each barcode were estimated. Results were also compared between various types of biological samples, i.e. eggs, infective larvae or adults.

Overall, the proposed COI barcode was suboptimal relative to the ITS-2 rDNA region, because of PCR amplification biases, a reduced sensitivity and higher divergence from the expected community composition. Metabarcoding yielded consistent community composition across the three sample types, although infective larvae may remain the most tractable in the field. Additional strategies to improve the COI barcode performances are discussed. These results underscore the critical need of mock communities for metabarcoding purposes.

## Introduction

Equine strongylids encompass a diverse fauna of 14 Strongylinae and 50 Cyathostominae described species (Lichtenfels et al., 2008). Among these, species from the genus *Strongylus* are responsible for the death of animals because of verminous arteritis liver pathology and peritonitis while Cyathostominae impinge on their host growth (McCraw and Slocombe, 1985, 1978, 1976; Reinemeyer and Nielsen, 2009). In addition, the mass emergence of developing cyathostomin stages can lead to a fatal syndrome of cyathostominosis characterized by abdominal pain, diarrhea or fever (Giles et al., 1985). The release of modern anthelmintics has drastically reduced the prevalence of Strongylus sp in the field as first mentioned in 1990 (Herd, 1990) and later confirmed by observations from necropsy data (Lyons et al., 2000; Sallé et al., 2020). However, treatment failure against cyathostomins has been found on many occasions across every continent for all drug classes currently available (Peregrine et al., 2014). Despite their worldwide distribution and relevance for stakeholders in the field, little knowledge has been gathered on the mechanisms driving their assemblage. Recent meta-analyses found that strongylid community structure was little affected by geo-climatic factors (Bellaw and Nielsen, 2020; Boisseau et al., 2021; Louro et al., 2021) and a few observations exist on the relationship between cyathostomin assemblage structure and environmental factors like temperature (Kuzmina et al., 2006; Nielsen et al., 2007), horse age (Bucknell et al., 1995; Kuzmina et al., 2016; Torbert et al., 1986) or the host sex (Kornaś et al., 2010; Sallé et al., 2018). The tedious and delicate process of species identification by morphological keys (Lichtenfels et al., 2008) is a major hurdle to study further the mechanisms of species assemblage, their turnover and the respective impacts of the host and their environment.

DNA-metabarcoding is a non-invasive, time- and cost-effective method for assessing nematode populations that provides data with comparable taxonomic resolution to morphological methods (Avramenko et al., 2015). This requires appropriate barcodes able to distinguish between the various phylogenetic strata. The internal transcribed spacer 2 region (ITS-2) of the nuclear rRNA cistron (Blouin, 2002; Kiontke et al., 2011) and the mitochondrial COI gene (Blaxter et al., 2005; Blouin, 2002; Prosser et al., 2013) have already been used for nematode molecular barcoding. For cyathostomin species, early barcoding attempts relied on the polymorphisms present in the ITS-2 rDNA region (Hung et al., 2000, 1999) before additional contributions were made using COI gene (McDonnell et al., 2000), or the longer intergenic spacer sequence (Cwiklinski et al., 2012). Additional work recently highlighted how the COI region could increase the resolution of species genetic diversity, suggesting a close phylogenetic relationship between *Coronocyclus coronatus* and *Cylicostephanus calicatus* (Bredtmann et al., 2019; Louro et al., 2021). In addition to this higher resolutive power, the protein-coding nature of the COI barcode can be leveraged to denoise sequencing data (Ramirez-Gonzalez et al., 2013). To date, metabarcoding experiments on equine strongylid species have exclusively focused on the ITS-2 rDNA cistron (Poissant et al., 2021). This may owe to the existence of universal primers and the length of the amplicon that is a good fit for short-read sequencing platforms. Observations in helminths also suggest that amplification efficiency is suboptimal for the COI region (Prosser et al., 2013) which speaks against its application for metabarcoding purposes. However, mitochondrial markers have better discriminating abilities between closely related or cryptic species (Bredtmann et al., 2019; Gao et al., 2020; Louro et al., 2021). Hence, the added value of the COI barcode remains to be determined for the study of cyathostomin species.

In addition, nemabiome approaches are biased in predicting relative taxon abundances (McLaren et al., 2019). These biases are inherent to various biological and technical factors including the DNA treatment procedures, the different number of cells represented by each taxon (that is tightly linked to the life-stage considered for strongylid species), PCR specifications (cycle number) and the amplification efficiency across taxa, and the genetic diversity (including structural variants and copy numbers) of the considered barcodes within taxa (Pollock et al., 2018). To date, the precision and recall of the metabarcoding approach applied to cyathostomins are unresolved. It is also currently unknown whether this approach provides a fair description of the actual species presence or absence, or their accurate relative abundances in their host. The impact of the considered life-stage, *i*.*e*. eggs, larvae or adult worms, has not been studied yet. Here, we aimed to address these three questions, i) the added value of the COI barcode to describe cyathostomin populations from mock samples in horses, ii) the predictive ability of the nemabiome approach to provide comparable results to morphological identification and iii) whether strongylid species can be correctly detected from different strongylid life-stages.

For this purpose, we developed degenerated primers to amplify the COI region following a strategy successful under other settings (Elbrecht et al., 2019; Elbrecht and Leese, 2017a), we built mock communities of diverse equine strongylid species and applied a nemabiome approach targeting the ITS-2 rDNA and COI gene regions. We compared the predictive performances of both approaches for various analytical pipelines and across diverse sample types and demonstrated that the ITS-2 rDNA barcode is a more reliable predictor of horses cyathostomin community taxonomical structure than the proposed COI barcode. Still, bioinformatic parameters need careful evaluation and amplification biases across species were found. For both barcodes, similar results were obtained from cyathostomin eggs, larvae or adult stages suggesting limited biases induced by the sample type. In conclusion, this study underscores the need for mock communities when studying equine strongylid communities with a metabarcoding approach.

## Materials and methods

### Mock community design and DNA extraction

Mock communities were built from morphologically identified equine strongylid specimens from pooled fecal samples in Ukraine (Kuzmina et al., 2016) and Poland. For each species, a single adult male was digested using proteinase K (Qiagen) in lysis buffer, before DNA extraction using a phenol/chloroform protocol. DNA was precipitated overnight in ethanol and sodium acetate (5M) at -20ºC and washed twice with 70% ethanol. The resulting DNA pellet was resuspended in 30 µL of TE buffer (10 mM Tris-HCl, 0.1 mM EDTA, pH 8.0) and DNa was quantified using using the Qubit^®^ double-stranded high sensitivity assay kit (Life Technologies™) with a minimal sensitivity of 0.1 ng/µL. Extracted DNA was stored at -20ºC. To quantify the impact of the community complexity, mock communities of 11 and five species were built and subjected to amplicon sequencing using the ITS-2 rDNA and mitochondrial COI barcodes (Table 1). Within each community, species DNA was either added on an equimolar basis or at their respective concentrations to mimic differences occurring in the field (heterogeneous communities; Table 1). Of note, the homogeneous communities comprised two *Cyathostomum pateratum* individuals to assess the impact of inter-individual variation. To further measure the resolution ability of the nemabiome approach, two-species communities were made with *Cyathostomum catinatum* and *C. pateratum*. Both species were either in imbalanced ratios (3-fold difference in both directions) or equal DNA concentration.

**Table 1.**
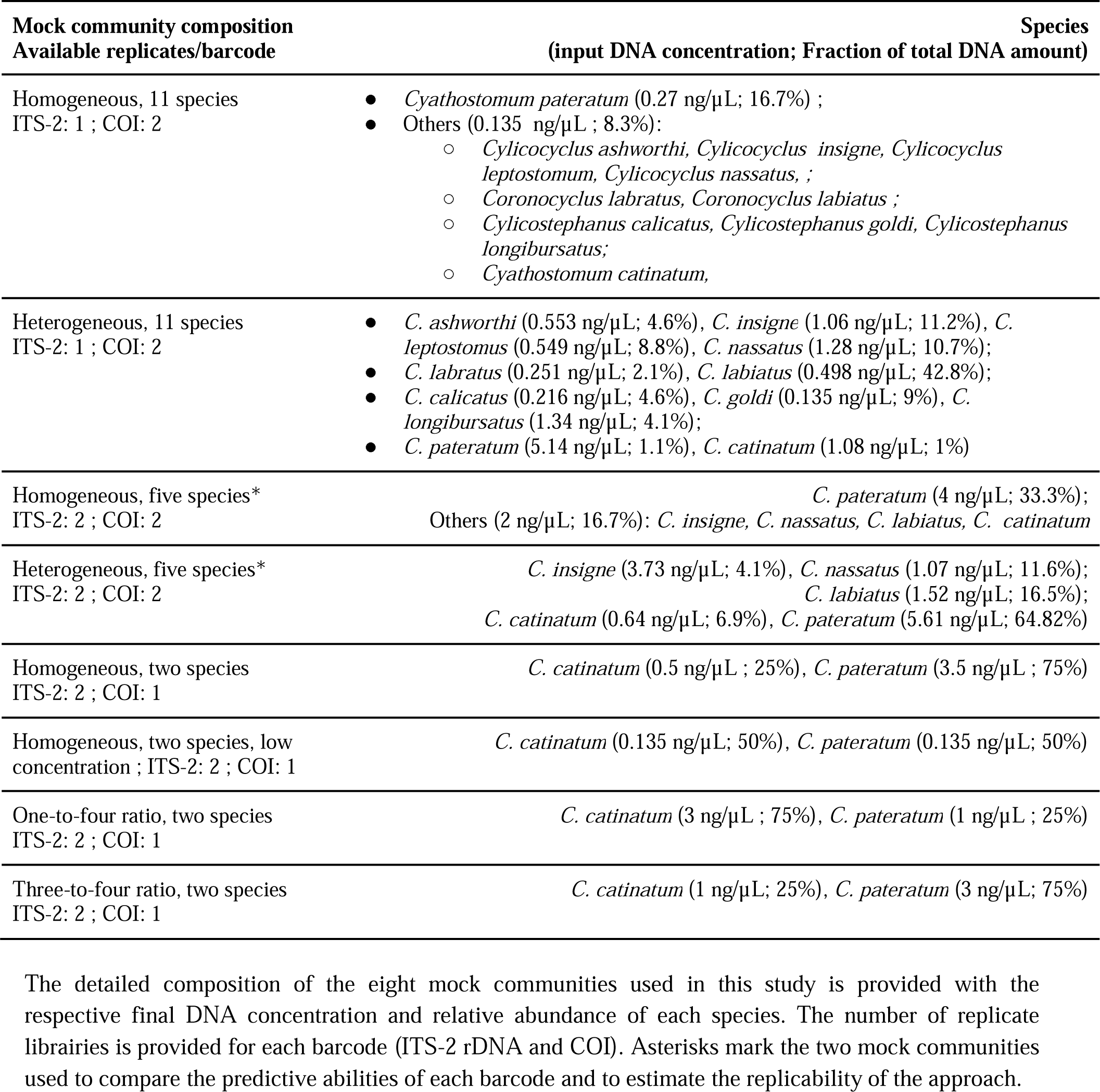
Detailed mock community composition

### Parasite material collection for comparison of the metabarcoding performances across sample type

Parasite material was collected from six Welsh ponies with patent strongylid infection. Faecal matter (200 g) was recovered from the rectum and incubated with 30% vermiculite at 25°C and 60% humidity for 12 days, before third-stage larvae were recovered using a Baerman apparatus. Strongylid eggs were extracted from another 200g of faeces. For this, faecal matter was placed onto a coarse sieve to remove large plant debris, before further filtering was made on finer sieves (150 µM and 20 µM mesh). Kaolin (Sigma K7375) was then added (0.5% w/v) to the egg suspension to further pellet contaminating debris (5 min centrifugation at 2,000 rpm). The supernatant was discarded and the egg pellet was resuspended in a dense salt solution (NaCl, d = 1.18) and centrifuged slowly (1,200 rpm for 5 min), before this final egg suspension was placed on a 20 µM mesh sieve for a last wash. Adult worms were collected from the same ponies at 18 and 21 hours after a pyrantel embonate treatment (Strongid⍰, Zoetis, France; 6.6 mg/Kg body weight). DNA extraction was performed as for the mock community samples. The amounts of adult worms, infective larvae and eggs are listed in Table 2.

**Table 2:**
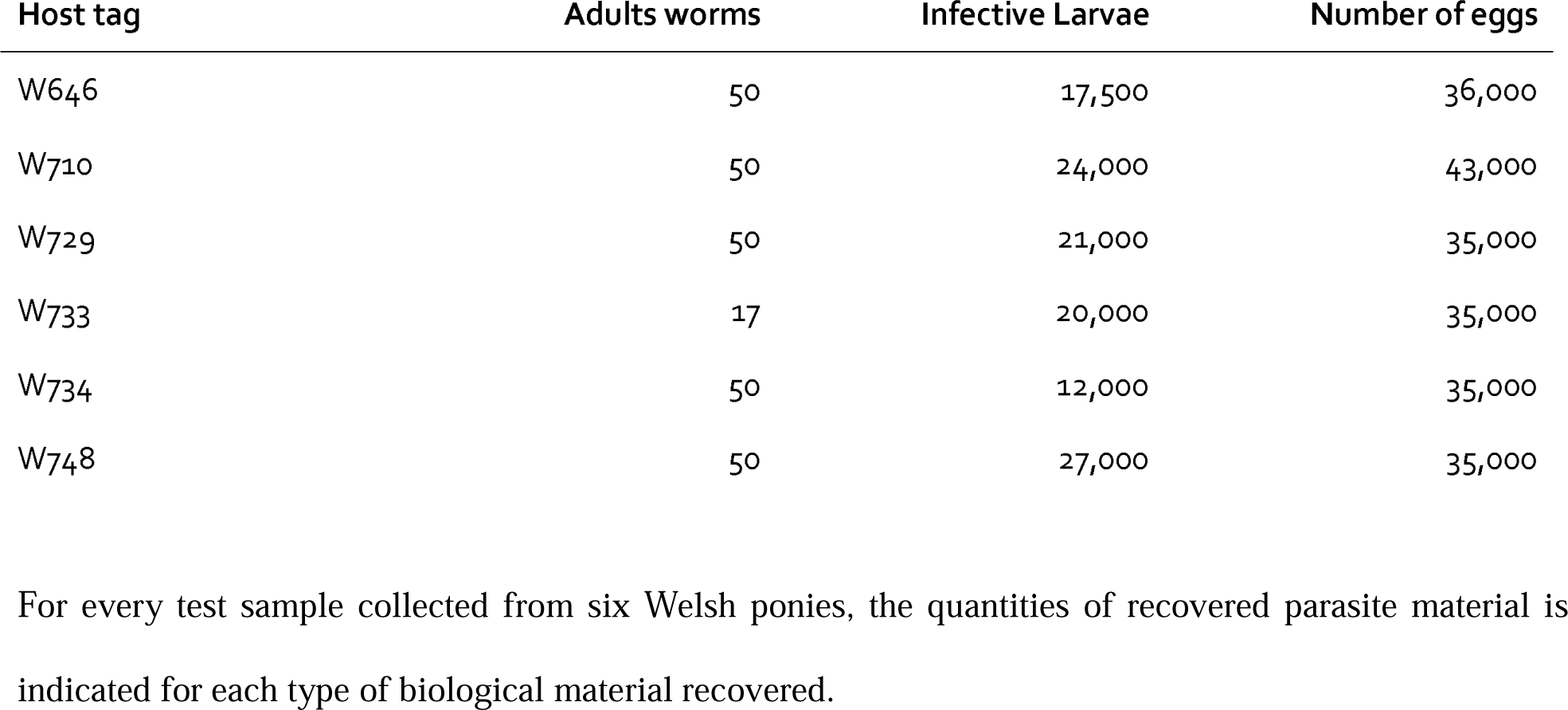
Quantity of larvae, adults and eggs used for DNA extractions. The quantities indicated are those present in each faecal aliquot.

| Host tag | Adults worms | Infective Larvae | Number of eggs |
| --- | --- | --- | --- |
| W646 | 50 | 17,500 | 36,000 |
| W710 | 50 | 24,000 | 43,000 |
| W729 | 50 | 21,000 | 35,000 |
| W733 | 17 | 20,000 | 35,000 |
| W734 | 50 | 12,000 | 35,000 |
| W748 | 50 | 27,000 | 35,000 |

For every test sample collected from six Welsh ponies, the quantities of recovered parasite material is indicated for each type of biological material recovered.

### COI and ITS-2 primer design

We aimed to define a 450-bp amplicon within the 650-bp fragment of the cytochrome c oxidase subunit I (COI) locus previously described (Bredtmann et al., 2019; Louro et al., 2021). This would leave at most 50 bp overlap, thereby allowing sequencing error correction and better amplicon resolution (Edgar and Flyvbjerg, 2015). For this, the mitochondrial sequences of 18 cyathostomin species with complete mitochondrial genomes available at that time (October 9th, 2020) were considered (AP017681: *Cylicostephanus goldi*, AP017698: *Strongylus vulgaris*; GQ888712: *Cylicocyclus insigne*; Q888717: *Strongylus vulgaris*; NC_026729: *Triodontophorus brevicauda*; NC_026868: *Strongylus equinus*; NC_031516: *Triodontophorus serratus*; NC_031517: *Triodontophorus nipponicus*; NC_032299: *Cylicocyclus nassatus*; NC_035003: *Cyathostomum catinatum*; NC_035004: *Cylicostephanus minutus*; NC_035005: *Poteriostomum imparidentatum*; NC_038070: *Cyathostomum pateratum*; NC_039643: *Cylicocyclus radiatus*; NC_042141: *Cylicodontophorus bicoronatus*; NC_042234: *Coronocyclus labiatus*; NC_043849: *Cylicocyclus auriculatus*; NC_046711: *Cylicocyclus ashworthi*). These sequences were aligned with Muscle v.3.8.21 (Edgar, 2004). Subsequently, this alignment was used to quantify sequence heterozygosity for 450-bp sliding windows using a custom python script (supplementary file S1). The consensus sequence of the region with the highest diversity, *i*.*e*. best discriminant across species, was isolated to design primers with the Primer3 blast web-based interface (Untergasser et al., 2012). Parameters were chosen to have an amplicon product of 400-450 bp, primers of 20 bp with melting temperatures of 60°C ± 1°C. Primer sequences were subsequently degenerated to account for identified SNPs in the mitochondrial sequence alignment, yielding a 24-bp long forward (5’ -RGCHAARCCNGGDYTRTTRYTDGG-3’) and 25-bp long reverse (5’ - GYTCYAAHGAAATHGAHCTHCTHCG-3’) primers. For the ITS-2 barcode, we relied on previously described primers (5’ -ACGTCTGGTTCAGGGTTGTT-3’) and NC2 (5’ - TTAGTTTCTTTTCCTCCGCT-3’) applied on cyathostomin communities already (Gasser et al., 1993). In both cases, a random single, double or triple nucleotide was added to the 5’ primer end to promote sequence complexity and avoid signal saturation. A 28-bp Illumina overhang was added for the forward and reverse sequences respectively, for subsequent ligation with Illumina adapters.

### Library preparation and sequencing

For library preparation, PCR reactions were carried out in 80 µl with 16 µl HF buffer 5X, 1.6 µl dNTPs (10mM), 4 µl primer mix containing forward and reverse primers, 0.8 µl Phusion High-Fidelity DNA Polymerase (2U/µl, Thermo Scientific), and 10 ng of genomic DNA. PCR conditions for ITS-2 were 95°C for 3 min for the first denaturation, then 30 cycles at 98°C for 15 s, 60°C for 15 s, 72°C for 15 s, followed by a final extension of 72°C for 2 min. For COI amplification, the PCR parameters were 95°C for 3 min, followed by a pre-amplification with 5 cycles of 98°C for 15 s, 45°C for 30 s, 72°C for 30 s, followed by 35 cycles of 98°C for 15 s, 55°C for 30 s, 72°C for 30 s then a final extension of 2 min at 72°C.

For each sample, 20 µl were examined on 1% agarose gel to check for the presence of a PCR amplification band at the expected product size (or absence thereof for negative controls). The concentrations of the purified amplicons were checked using a NanoDrop 8000 spectrophotometer (Thermo Fisher Scientific, Waltham, USA), and the quality of a set of amplicons was checked using DNA 7500 chips onto a Bioanalyzer 2100 (Agilent Technologies, Santa Clara, CA, USA). A homemade six-bp index was added to the reverse primer during a second PCR with 12 cycles using forward primer (AATGATACGGCGACCACCGAGATCTACACTCTTTCCCTACACGAC) and reverse primer (CAAGCAGAAGACGGCATACGAGAT-index-GTGACTGGAGTTCAGACGTGT) for single multiplexing. The final libraries had a diluted concentration of 5 nM to 20 nM and were used for sequencing. Amplicon libraries were mixed with 15% PhiX control for quality check. Libraries were further processed for a single run of MiSeq sequencing using the 500-cycle reagent kit v3 (2 × 250 output; Illumina, USA).

### Analytical pipelines

#### Quality control and filtering

Sequencing data were first filtered using cutadapt v1.14 to remove insufficient quality data (-q 15), trim primer sequences, and remove sequences with evidence of indels (--no-indels) or that showing no trace of primer sequence (--discard-untrimmed).

#### Analytical pipelines for community structure inference using the ITS-2 amplicon

The implemented framework was similar to previous work (Poissant et al., 2021) that used the DADA2 algorithm (Callahan et al., 2016) to identify amplicon sequencing variants after error rate learning and correction. The denoising procedure, consisting in learning error rates independently for both forward and reverse reads, was applied for two discrete stringency parameter sets either tolerating a single error for both reads (mxEE = 1) or more relaxed stringency (mxEE = 2 and 5 for the forward and reverse reads respectively). Truncation length of forward and reverse reads was set to 200 or 217 bp. Last, the band_size parameter effect was also explored considering three values, i.e. -1, 16 and 32, that respectively disable banding and implement the default or a more relaxed value that is recommended for ITS-2 sequences. In every case, denoising was run using the pseudo-pool option and chimera detection relied on the consensus mode.

Taxonomic assignment was subsequently performed using a sequence composition approach using the IDTAXA algorithm (Murali et al., 2018) as implemented in the DECIPHER R package v.2.18.1 with minimal bootstrap support of 50%. This last step relied on the curated ITS-2 rDNA database for nematodes (Workentine et al., 2020), last accessed on February 3rd 2022.

#### Analytical pipelines for community structure inference using the COI amplicon

For the COI barcode, we built a custom COI barcode sequence database for Cyathostominae and Strongylidae species collected from Genbank, BOLD database using the Primerminer package v.0.18 (Elbrecht and Leese, 2017b) and concatenated into a single fasta file. Sequences were subsequently edited to remove elephant Cyathostominae species (*Quilonia* sp, *Murshidia* sp, *Kilonia* sp, and *Milulima* sp) using the seqtk v.1.3 subseq option (https://github.com/lh3/seqtk). Some entry names (n = 18) consisted of an accession number that was manually back-transformed to the corresponding species name. Duplicate entries were removed with the rmdup option of the seqkit software v.0.16.0 (Shen et al., 2016) and sequences were dereplicated using the usearch v11.0.667 -fastx_uniques option (Edgar, 2010). To reduce the database complexity and promote primary alignment, the most representative sequences were further determined using the usearch -cluster_smallmem option, considering two identity thresholds of 97 and 99%. Amplicon analysis relied on a mapping approach to the custom COI sequence database implemented using the minimap2 software v.2-2.11 (Li, 2018) as described elsewhere (Ji et al., 2020). First, paired-end reads were merged into amplicon sequences using the usearch software v11.0.667 and the -fastq_mergepair option (Edgar, 2010). Merged amplicon sequences were subsequently mapped onto the COI sequence database using the minimap2 short read mode (-ax sr) (Li, 2018). Mapping stringency was varied to select the most appropriate combinations using k-mer sizes of 10, 13, and 15 (default), window sizes w of 8, 9 or 10 (default), and varying mismatch penalty values (B = 1, 2, 3 or the default values 4). The lower the value, the more permissive the alignment for these three parameters. Produced alignments were converted to bam files using samtools v1-10 after filtering against unmapped reads, alignments that were not primary and supplementary secondary alignments using the -F 2308 flag (Li et al., 2009). To evaluate how mapping stringency, filtered bam files were also produced using a mapping quality cut-off of 30. Species abundance was then inferred from read depth over each COI sequence that was determined using the bedtools genomecov algorithm (Quinlan and Hall, 2010) and scaled by the sequence length.

### Quantitative PCR (qPCR) assay for species-specific amplification of the ITS-2 rDNA region

To quantify any biases in PCR amplification, single-species DNA used to make mock communities was subjected to quantitative PCR reactions with the ITS-2 and COI-specific primers. The DNA was diluted at 1:250 and 1□µl of DNA was added to each reaction. qPCRs were carried out on a Biorad CFX Connect Real-Time PCR Detection System following the iQ SYBR GREEN supermix® protocol (Biorad, France, 1708882). Reactions were run in triplicate for each species with 40 amplification cycles: 95°C for 3 min for the first denaturation, then 45 cycles of 98°C for 15 s, 60°C for 30 s, 72°C for 40 s, followed by a melt curve (65°C to 95°C).

### Statistical analyses

Statistical analyses were run with R v.4.0.2 (R Core Team, 2016). Community compositional data were imported and handled with the phyloseq package v1-34.0 (McMurdie and Holmes, 2013). Abundance data (read count for the ITS-2 rDNA region, or scaled read depth for COI) were aggregated at the species level using the *taxglom()* function of the phyloseq package v1-34.0 and converted to relative abundances for further analysis. These data were used to compare the predictive ability of each barcode and pipeline in a first comparison. After the best bioinformatics parameters were identified for each barcode, the ASV count tables were further filtered to remove putative contaminants or spurious signals representing either less than 100 reads per base pair for the COI barcode or 40 reads for the ITS-2 rDNA barcodes.

Species richness, alpha-diversity and beta-diversity analyses were conducted with the vegan package v2.5-7 (Oksanen et al., 2017). PerMANOVA was implemented using the *adonis()* function of the vegan packagev2.5-7 (Oksanen et al., 2017).

To monitor the predictive ability of each pipeline and barcode, the precision (the proportion of true positives among all positives called, i.e. true positives and false positives) and recall (the proportion of true positives among all true positives, i.e. true positives and false negatives) of the derived community species composition were computed and combined into the F1-score as:

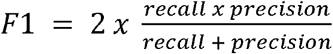

This score supports the ability of a method to correctly identify species presence while minimizing the number of false-positive predictions (trade-off between precision and recall) (Hleap et al., 2021).

Alpha diversity was estimated using the Shannon and Simpson’ s indices using the *estimate_richness()* function of the phyloseq package (McMurdie and Holmes, 2013). To compare between conditions and barcodes, the difference between the mock community expected and observed alpha diversity values was further considered. The divergence between the inferred community species composition and the true mock community composition was estimated from a between-community distance matrix based on species presence/absence (Jaccard distance) or species relative abundances (Bray-Curtis dissimilarity) using the *vegdist()* function of the vegan package (Oksanen et al., 2017). Differences in nemabiome between sample types were visualized with a non-metric multidimensional scaling (NMDS) with two dimensions using Jaccard and Bray-Curtis dissimilarity.

For each of these variables (F1-score, alpha-diversity differences and divergence) and within each barcode, estimated values were regressed upon bioinformatics pipeline parameters (mxee, truncation length and band size parameters for the ITS-2 barcode data; k-mer size, window size, mismatch penalty and mapping quality for the COI barcode) to estimate the relative contribution of these parameters and determine the most appropriate combination for each barcode independently. The model with the lowest Akaike Information Criterion value was first selected with the *stepAIC*() function of the R MASS package v.7.3-55 (Venables and Ripley, 2002) to retain the most relevant parameter combination (model with the lowest AIC). Parameter values were then chosen according to their least-square mean estimate. These analyses were applied to every available data within each barcode.

These variables were subsequently compared between the ITS-2 and COI barcodes using the best bioinformatic pipeline and across every community available for each barcode. Replicability was estimated from the two mock communities with five species, available for both barcodes (Table 1) and the contribution of species and replicate run effects were determined using the same AIC-based procedure.

To estimate amplification efficiencies, the threshold cycle (Ct) values were regressed upon the log10-transformed DNA concentration for each species and barcode. The PCR efficiency was subsequently derived as: 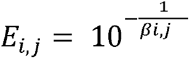, where *E*_*i,j*_ is the efficiency, and *βi, j* stands for the regression slope of species *i* and barcode *j*. The R script and the necessary files used for analysis are available under the github repository at https://github.com/guiSalle/Horse_Nemabiome_benchmark. The associated raw sequencing will be made available upon manuscript acceptance on the SRA platform.

## Results

### Impact of the community size on the ITS-2 and COI barcode performances

The best bioinformatic parameters were determined for each barcode according to their predictive performances of mock community composition (Supplementary information and supplementary Tables 1, 2 and 3). This comparison relied on the same two mock communities of five species (Table 1). The combination of the minimap default values for the mismatch penalty (B = 4) and the window size (w = 10) parameters, with a k-mer size of 10 base pairs and no further filtering on mapping quality (MQ = 0) was deemed as the most optimal pipeline for the COI barcode. For ITS-2, stringent tolerance in the maximal expected number of errors and a truncation length of 200 bp was the best parameters for cyathostomin community structure prediction.

With these settings, the number of reads available for the considered mock communities ranged between 6,164 and 151,606 reads for the COI barcode, while these numbers ranged between 816 and 86,424 non-chimeric sequences for the ITS-2 rDNA barcode.

No significant difference was found in the F1 score between the ITS-2 and COI barcodes (*F*_*1,19*_ = 0.26, *P* = 0.61; Figure 1). In terms of species detection, the COI barcode ability to recover the true species composition was suboptimal: this barcode recovered nine correct species at most in the most complex communities and systematically overlooked *C. leptostomum* and *C. labratus* in these. On the contrary, it also identified *C. coronatus* (in all of the four 11-species mock communities) and *C. minutus* (in one out of the four 11-species mock communities) despite these species not being present. Their relative abundances remained however lower than 1% for *C. coronatus* and 0.02% for *C. minutus* and was associated with low mapping quality (Phred quality score < 6). The ITS-2 rDNA barcode performed better with ten species detected overall but *C. goldi* was systematically overlooked in the 11-species mock communities.

**Figure 1.**
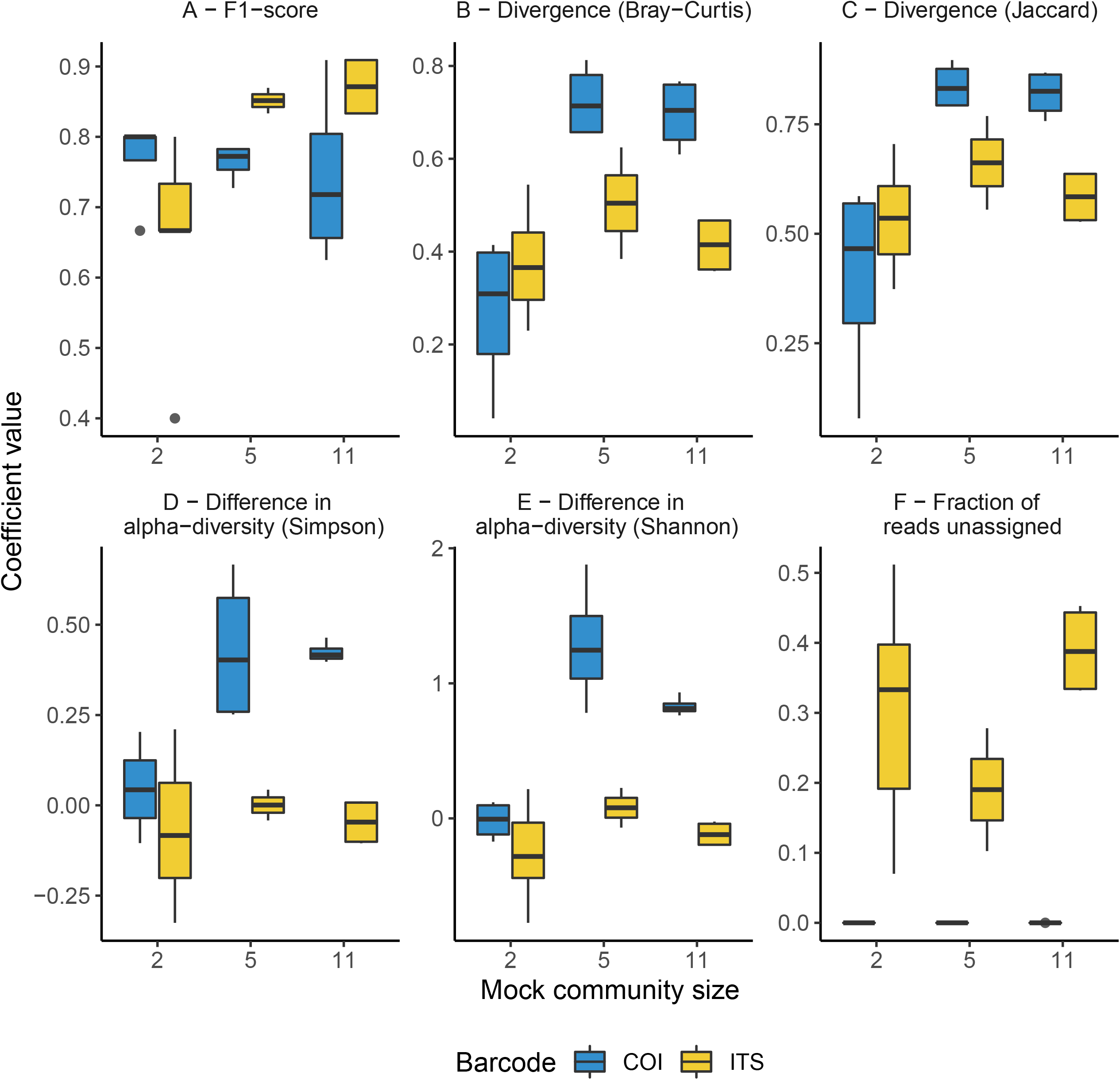
Comparison of the predictive abilities of cyathostomin community structure for the mitochondrial COI and ITS-2 rDNA barcodes. Considered coefficient values are represented across three mock community sizes for the mitochondrial COI (blue) and ITS-2 rDNA (yellow) barcodes. F1-score corresponds to the trade-off between identifying true positives while minimizing the false discovery rate (panel A). Divergence was computed as the Bray-Curtis (species relative abundances; panel B) or Jaccard distances (species presence/absence; panel C) between the expected and observed mock community composition. Differences between observed and expected alpha diversity are given in panels D and E. Panel F depicts the fraction of reads with no taxonomy assigned.

The ITS-2 barcode gave better representations of the most complex cyathostomin communities composition (with five or eleven species; Figure 1B, C). for species relative abundances (reduction of 0.14 ± 0.07 in Bray-Curtis dissimilarity relative to COI, *P* = 0.04) or species presence/absence (decrease of 0.18 ± 0.08 in Jaccard distance relative to COI, *P* = 0.04). The ITS-2 barcode also gave the closest estimates of the expected alpha diversity (*F*_*1,19*_ = 22.9, *P* = 1.2 × 10^−4^ and F_1,19_ = 34.9, *P* < 10^−4^ for Simpson and Shannon indices; Figure 1). These differences vanished however when considering the mock communities composed of two *Cyathostomum* species (Figure 1): no statistical difference was found in any of the six parameters in that case (*P* > 0.2 in all cases).

Last, the fraction of unassigned reads increased with the mock community size for the ITS-2 rDNA barcode (ranged between 17% to 35%, Figure 1) but remained negligible with the COI region as a result of the mapping procedure (2 × 10^−5^ % for the most complex community and null otherwise; Figure 1).

In summary, none of the two barcodes offered a perfect fit for the expected community composition, but the ITS-2 rDNA barcode was closer to the truth.

### Replicability and correspondence between read counts and input DNA for the ITS-2 and COI barcodes

Pearsons’ correlation between log-transformed species abundance measured across two runs for two 5-species mock communities (Table 1) was 98% for both barcodes. In line with this consistency across libraries, input DNA concentration modeling performed best when using the observed read counts and species as predictors (model AIC = 15.8 and 27.9; *R*^*2*^ = 87.3% and 73.9% for ITS-2 and COI barcodes). The replicate effect was negligible in both cases (*F*_*1,17*_ = 0.2, *P* = 0.66 and *F*_*1,17*_ = 0.001, *P* = 0.99 for the ITS-2 and COI respectively; Figure 2).

**Figure 2.**
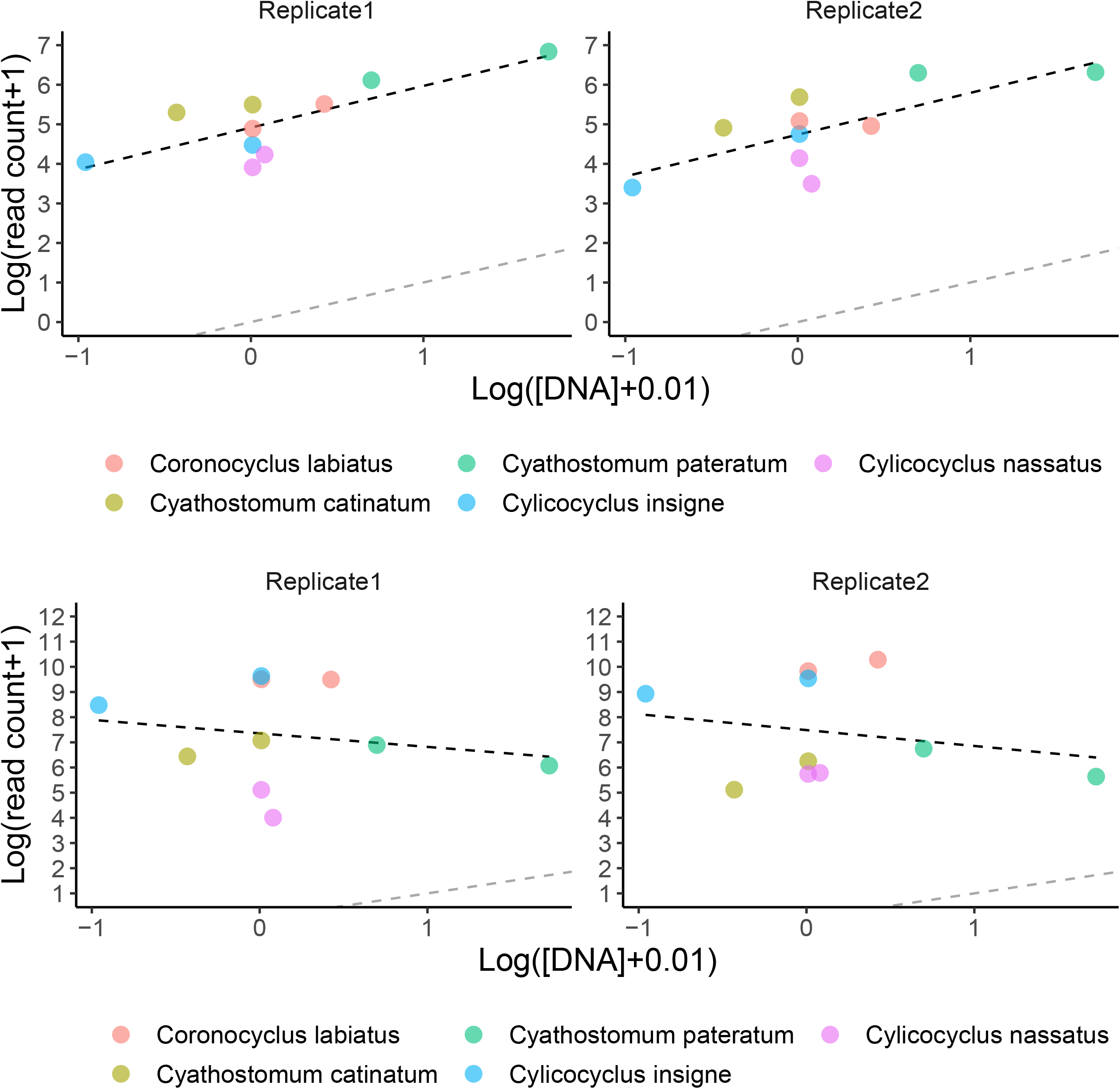
The relationship between input DNA concentration and observed counts across two runs. Recovered species counts for the ITS-2 rDNA (top panels) and COI (bottom row) barcodes are plotted against the respective input DNA concentration before amplification for two replicate samples (left and right panels). Regression slopes of read count upon DNA concentration are represented (black dashed line) and the grey dashed line would fit perfect correlation between input DNA and observed read counts.

Despite similar behaviours in terms of modeling species DNA abundance, the correlation between observed read counts and species input DNA quantity was 75% for the ITS-2 rDNA barcode but non-significantly different from 0 for the COI barcode (Pearson’ s *r* = -0.21, *P* = 0.38; Figure 2).

Hence, the designed COI barcode appeared suboptimal in its ability to provide a fair representation of the input cyathostomin DNA abundance, in sharp contrast with the ITS-2 barcode.

### PCR amplification bias for the COI and ITS-2 rDNA barcodes

The imperfect correlations found between input DNA quantity and observed read counts might relate to biases in the first PCR amplification. To test this hypothesis, qPCRs were performed on each species DNA from the same single individual used for library preparation (Table 3). The average amplification efficiency was 67.7% ± 0.24% for the COI barcode. It was above 90% for the two *Cyathostomum* species (Table 3) but it fell below 70% for five species. Among these, *C. calicatus* and *C. longibursatus* showed the lowest values (39% and 13% respectively, Table 3). Omitting the outlier values found for the *Cylicostephanus* members (*C. goldi* undetected and too high efficiency for *C. calicatus*; Table 3), the ITS-2 rDNA barcode yielded more consistent and higher amplification efficiency on average (92.2% ± 0.03%; *t*_*10*_ = 3.42, *P* = 0.006) than COI.

**Table 3.** Species-specific amplification efficiencies of the COI and ITS-2 barcodes for 11 cyathostomin species

| Barcode | Genus | Species | Slope | Std. error | Intercept | Efficiency | R<sup>2</sup> |
| --- | --- | --- | --- | --- | --- | --- | --- |
| COI | *Coronocyclus* | *C. labratus* | -3.90 | 0.52 | -1.48 | 0.81 | 0.87 |
|  |  | *C. labiatus* | -4.70 | 0.44 | -5.12 | 0.63 | 0.93 |
|  | *Cyathostomum* | *C. pateratum* | -3.45 | 0.55 | 1.90 | 0.95 | 0.82 |
|  |  | *C. catinatum* | -3.53 | 0.22 | 2.70 | 0.92 | 0.97 |
|  | *Cylicocyclus* | *C. leptostomum* | -3.98 | 0.45 | -3.93 | 0.78 | 0.91 |
|  |  | *C. ashworthi* | -4.34 | 0.35 | -3.92 | 0.70 | 0.95 |
|  |  | *C. nassatus* | -4.36 | 0.47 | -5.32 | 0.70 | 0.91 |
|  |  | *C. insigne* | -4.38 | 0.39 | -5.30 | 0.69 | 0.94 |
|  |  | *C. calicatus* | -6.99 | 0.71 | 20.89 | 0.39 | 0.92 |
|  | *Cylicostephanus* | *C. goldi* | -3.98 | 1.91 | 29.73 | 0.78 | 0.29 |
|  |  | *C. longibursatus* | -19.06 | 3.77 | -7.28 | 0.13 | 0.89 |
| ITS-2 | *Coronocyclus* | *C. labratus* | -3.48 | 0.11 | 14.47 | 0.94 | 0.99 |
|  |  | *C. labiatus* | -3.53 | 0.13 | 15.93 | 0.92 | 0.99 |
|  | *Cyathostomum* | *C. pateratum* | -3.33 | 0.13 | 15.91 | 1.00 | 0.99 |
|  |  | *C. catinatum* | -3.64 | 0.14 | 15.18 | 0.88 | 0.99 |
|  | *Cylicocyclus* | *C. leptostomum* | -3.47 | 0.07 | 14.24 | 0.94 | 1.00 |
|  |  | *C. ashworthi* | -3.49 | 0.13 | 15.63 | 0.94 | 0.99 |
|  |  | *C. nassatus* | -3.43 | 0.11 | 15.57 | 0.95 | 0.99 |
|  |  | *C. insigne* | -3.43 | 0.12 | 15.36 | 0.96 | 0.99 |
|  |  | *C. calicatus* | -3.64 | 0.08 | 14.41 | 0.88 | 1.00 |
|  | *Cylicostephanus* | *C. goldi* | 0.00 | 0.40 | 33.09 | -1.00 | -0.06 |
|  |  | *C. longibursatus* | -0.98 | 0.88 | 26.65 | 9.53 | 0.03 |

Regression coefficients (slope, associated standard error and intercept) of the amplification efficiency (measured with Ct) upon DNA concentration are given for every cyathostomin species considered. Derived amplification efficiency and regression explanatory power are also provided. Data are ordered by genus and efficiency.

### The considered life stage has limited effect on the inference of community composition

Nemabiome data were generated from six horses for three life stages (eggs, infective larvae and adult worms collected after pyrantel treatment). The total number of species present in this population remains unknown to date. The COI barcode retrieved 12 species, of which *C. radiatus* was not found with the ITS-2 barcode. This latter barcode allowed the retrieval of 13 species and four additional amplicon sequence variants that were assigned at the genus level only (Figure 3). ITS-2 data also recovered *C. leptostomum* and *Craterostomum acuticaudatum* that were not found with the COI marker (Figure 3).

**Figure 3.**
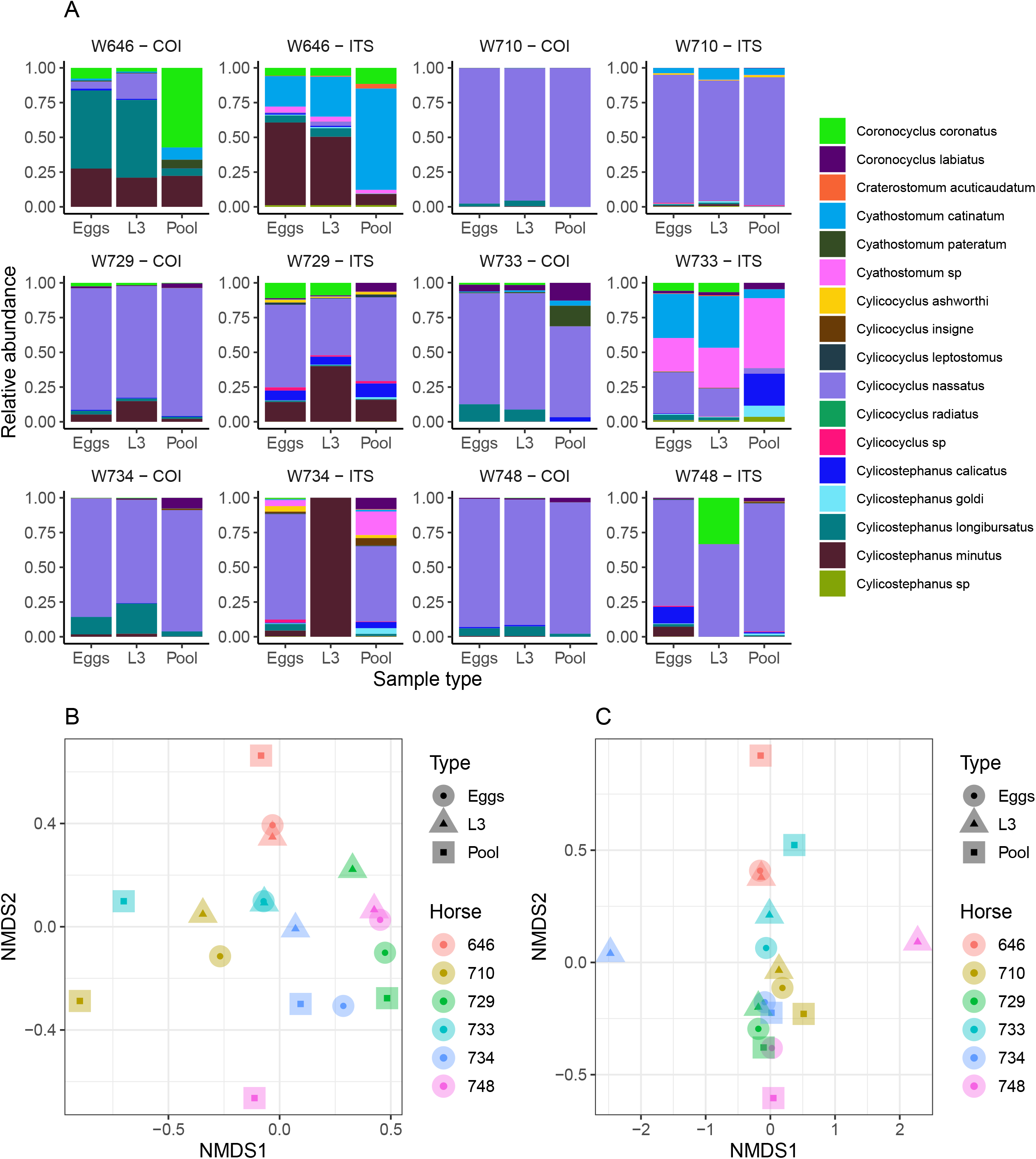
Impact of the considered life-stage on predicted equine strongylid community diversity. Panel A provides the relative abundances measured in the strongylid communities from six horses using the COI or ITS-2 barcodes and across three sample types. Panels B and C show the first two axes of a Non-linear Multidimensional Scaling analysis based on Bray-Curtis dissimilarity for the COI and ITS-2 rDNA region respectively. The plot illustrates the consistency of species relative abundances across sample types and the reduced alpha diversity for the COI barcode.

The Shannon index showed no significant variation across the considered life-stages for both barcodes (*P* = 0.96, Table 4) as the observed differences fell below the resolutive power of this experiment (difference of 1.8 detectable with a significance level of 5% and power of 80%).

**Table 4.** Shannon index estimated using three sample types.

| Type | Barcode | Average ± standard error |
| --- | --- | --- |
| Eggs | COI | $0.58 \pm 0.36$ |
| L3 | COI | $0.65 \pm 0.32$ |
| Pool | COI | $0.58 \pm 0.46$ |
| Eggs | ITS-2 | $1.11 \pm 0.47$ |
| L3 | ITS-2 | $0.93 \pm 0.66$ |
| Pool | ITS-2 | $1.10 \pm 0.62$ |

For each sample type, the average Shannon index estimated across six replicates and associated standard error is provided.

Alpha-diversity estimates obtained with the ITS-2 barcode were higher (difference of 0.44 ± 0.16 relative to COI, *t*_*1*_ = 2.75, *P* = 0.01; Table 4).

The community structure was generally in good agreement across sample types for both the ITS-2 and COI barcodes, although two larval samples exhibited outlying behaviours with the ITS-2 barcode (Figure 3). In agreement with this observation, PerMANOVA showed that the sample type was not a significant driver of the beta-diversity, explaining 16.8% (*P* = 0.12, *F*_*2,15*_ = 1.51, *R*^*2*^ = 0.168) and 9.9% (*P* = 0.63, *F*_*2,15*_ = 0.83, *R*^*2*^ = 0.099) of the variance in species relative abundances for the COI and ITS-2 barcode respectively.

Altogether, these observations suggest that metabarcoding would yield the same results from cyathostomin eggs extracted from the faecal matter, infective larvae harvested after egg culture or adult worm collection after treatment.

## Discussion

This study proposed a new barcode based on the COI gene and quantified the effect of a few factors affecting the metabarcoding of horse strongylid communities, including bioinformatic pipeline parameters, the number of species in the community, the barcode amplification step or the considered sample type. Overall, the proposed COI barcode appears suboptimal in comparison to the ITS-2. The use of a mock community is of utmost importance to fine tune the bioinformatic pipeline parameters while the approach appears to be robust across the various sample types available.

The COI region has been used for nemabiome or phylogenetic studies of other nematode species, including *Haemonchus contortus* (Blouin, 2002) or some free-living *Caenorhabditis* species (Kiontke et al., 2011). Its higher genetic variation poses this marker as an ideal barcode for cyathostomin species with evidence that it could better delineate the phylogenetic relationship between *Coronocyclus coronatus* and *Cylicostephanus calicatus* members (Louro et al. 2021). This study aimed to produce amplicons suitable for merging of read pairs, i.e. with a total length lower than 500 bp. To compensate for the diversity found across species over that region, primer sequences were degenerated and a 5-cycle at lower temperature pre-amplification was applied to compensate for suboptimal priming (Kwok et al., 1994) while subsequent mapping stringency was lowered to promote read mapping against the reference sequence database. As can be expected, this strategy yielded a poor correlation between input DNA and quantified reads. Specifically, the PCR amplification efficiency showed significant variation across species with suboptimal amplification for *C. calicatus* and *C. longibursatus* species. Additional factors such as the type of DNA polymerase could also affect the results and warrant further investigations (Nichols et al., 2018). The designed COI barcode did not outperform the ITS-2 rDNA region in terms of species detection or Jaccard-based (species presence or absence) diversity estimates. In this respect, inclusion of *C. coronatus* in mock communities may have yielded less favourable results for the ITS-2 rDNA barcode as this species entertains close phylogenetic relationship with *C. calicatus* (Bredtmann et al., 2019).

While the described approach was suboptimal, other strategies targeting the mitochondrial genome could still be applied (Ji et al., 2020; Liu et al., 2016). First, bulk shotgun sequencing of cyathostomin populations could be used for mapping against a reference database of mitogenomes as applied to arthropods (Ji et al., 2020). This strategy improved the correlation between species input DNA and the number of mapped reads (Ji et al., 2020). This is however more expensive and is limited by the available mitogenomic resources for equine strongylids (18 species available at the time of this experiment). Primer cocktail to simultaneously amplify multiple amplicons is another alternative that may increase the range of diversity being sampled (Chase and Fay, 2009). However, its performance under other settings was poor and was not better than relying on degenerated primers (Elbrecht et al., 2019). Last, third-generation sequencing technologies like the Pacific Biosciences and Oxford Nanopore Technologies which are both able to sequence long DNA fragments, could recover the whole COI gene or the mitochondrial genome from a pool of strongylid species. The portable Oxford Nanopore Technologies device offers a convenient set-up that can be deployed in the field (Quick et al., 2015), and could deliver full-length COI barcode data for up to 500 insect specimens in a single run (Srivathsan et al., 2018). This comes, however, at the cost of sequencing errors associated with insertion/deletion errors over homopolymeric regions (Srivathsan et al., 2018). But this drawback can be overcome as the protein-coding nature of the COI gene provides a solid basis for error denoising (Andújar et al., 2018; Ramirez-Gonzalez et al., 2013).

The ITS-2 rDNA region has been implemented already for characterizing equine strongylid communities (Poissant et al., 2021) but its predictive performances have not been characterized against mock equine strongylid communities. Its amplification was more robust as already reported (Louro et al., 2021) and the PCR amplification efficiency was consistent across considered species. The lower amplification found for the two *Cylicostephanus* species mirrored that found with COI and may relate to the input material. It remains unclear however why the recorded amplifications were slightly lower for *C. catinatum* and *C. calicatus*. The presence of polymorphisms in the latter species has been described and could have hampered PCR amplification (Louro et al., 2021).

Despite the breadth of species considered in this work, it was not possible to cover every known member of the equine Cyathostominae and Strongylidae families. Past investigation has focused on *Strongylus* spp showing that the metabarcoding was underestimating their true abundances (Poissant et al., 2021). Among the Cyathostominae family, species of intermediate abundance and prevalence like *C. coronatus* and *C. radiatus* should be considered for further studies. Specifically, the ability of the various approaches and algorithms to delineate between *C. calicatus* and *C. coronatus* members should be further investigated. As our observations suggest that pipeline performances were dependent on the species number, an investigation of more complex communities remains to be completed.

In turn, nemabiome is expected to unravel yet unknown facets of cyathostomin phenology, like their seasonal preference, or any priority effects between species (Boisseau et al., 2021). The lack of significant differences between the considered sample types for this approach supports a flexible implementation in the field. Infective larvae certainly can be harvested a few days after sample collection and as such, they remain the most convenient sample type for field work. In contrast, egg samples will develop into first-stage larvae within 24-48 hours and adult collection will be dependent on the species drug sensitivity.

To overcome the described challenges owing to barcode properties, metagenomic shotgun sequencing on DNA extracted from faeces could resolve the horse gut biodiversity in a single-pot experiment. While gut microbial gene catalogs have been built recently (Ang et al., 2022; Mach et al., 2022), the genomic resources for cyathostomin remain restricted to a few mitochondrial genomes that span the *Coronocyclus* (Yang et al., 2020), *Cyathostomum* (Wang et al., 2020), *Cylicocyclus* (Y. Gao et al., 2017), *Cylicostephanus* (Gao et al., 2020), and *Triodontophorus* (J.-F. Gao et al., 2017) genera, and a single heavily fragmented genome assembly for *C. goldi* (International Helminth Genomes, 2019).

## Conclusion

This work investigated the impact of experimental factors on metabarcoding approaches applied to equine strongylids. Overall, the developed COI barcode was suboptimal relative to the ITS-2 region. But this latter barcode displayed variable PCR amplification efficiency that may bias subsequent inferences. Cyathostomin larvae appear to be the most valuable biological material for metabarcoding. However, reliance on eggs extracted from the faecal matter or adult worms yielded the same results and could be considered for studies.

Metagenomic sequencing should offer a viable option to overcome the limitations of barcodes but is faced with limited available genomic resources for the species of interest.

## Acknowledgements

The authors are grateful to the INRAE UEPAO equine facility for their support in implementing this experiment. We are grateful to the genotoul bioinformatics platform Toulouse Occitanie (Bioinfo Genotoul, https://doi.org/10.15454/1.5572369328961167E12) for providing computing and storage resources.

