## Supplementary information for "Evaluation of the nemabiome approach for the study of equine strongylid communities"

### 1. **Impact of dada2 parameters on the ITS-2 taxonomy assignment and community composition inference**

To determine the best combination of dada2 parameters (supplementary Table 1), the trade-offs between recall and precision (F1-score), divergence from the expected community structure and alpha-diversity were estimated (supplementary Table 2). The shorter truncation length (200 bp) performed better than for 217 bp (0.1 difference in F1-score on average, *P* = 4.6 x 10^-5^). With this truncation length, the best F1-score was obtained when the most stringent maximal expected number of errors (mxee parameter) was applied (supplementary Table 3). But more relaxed parameters slightly reduced the divergence between the predicted community composition and the expected mock structure (supplementary Table 2), although this was not true for the most complex communities (supplementary Table 2).

The band size parameter had negligible effect on the F1-score and alpha-diversity measures (supplementary Table 2), but the ITS-2 recommended value yielded community composition closer to the expected values both in terms of species relative abundance and species presence/absence (supplementary Table 2). The taxonomy assignment missing rate was also better for this parameter value ( 10.8 % ± 2.8% reduction relative to a band size value of -1, *P* < 10^-4^) and was considered for further analysis.

Overall, stringent tolerance in the maximal expected number of errors and a truncation length of 200 bp was the best parameters for cyathostomin community structure prediction.

**Supplementary Table 3. Effect of bioinformatic parameters on community composition for the ITS-2 and COI barcodes**

|  | **Parameter** | **Level** | **F1** | **Distance from truth**  **(Bray-Curtis)** | **Distance from truth**  **(Jaccard)** | **Difference in ɑ-diversity (Shannon)** | **Difference in ɑ-diversity (Simpson)** |
| --- | --- | --- | --- | --- | --- | --- | --- |
| ITS | Maximal expected error | mxee = 1,1 | *ref. level* | *ref. level* | *ref. level* | *ref. level* | *ref. level* |
|  |  | mxee = 2, 5 | **0.06 ± 0.03** | **-0.12 ± 0.03** | **-0.11 ± 0.03** | 0.07 ±0.04 | 0.03 ±0.02 |
|  | Truncation length | 200 bp | *ref. level* | *ref. level* | *ref. level* | dAIC = 0.01 | dAIC = 8 x 10-4 |
|  |  | 217 bp | **-0.09 ± 0.03** | 0.13 ± 0.03 | 0.12 ± 0.03 |  |  |
|  | Band size | BS = -1 | dAIC = 0.01 | *ref. level* | *ref. level* | dAIC = 0.18 | *ref. level* |
|  |  | BS = 16 |  | 0.002 ± 0.03 | 0.002 ± 0.03 |  | -0.002 ± 0.03 |
|  |  | BS = 32 |  | **-0.09 ± 0.03** | **-0.09 ± 0.03** |  | 0.05 ± 0.03 |
| COI | Mismatch penalty | B = 1,2,4 | dAIC = 0.06 | dAIC = 0.03 | dAIC = 0.01 | dAIC = 0.06 | dAIC = 0.05 |
|  | k-mer size | k = 10 | ref. level | dAIC = 0.06 | dAIC = 0.03 | dAIC = 0.17 | dAIC = 0.05 |
|  |  | k = 13 | -0.02 ± .02 |  |  |  |  |
|  |  | k = 15 | **-0.05 ± 0.02** |  |  |  |  |
|  | Window size | w = 8 | dAIC = 0.03 | dAIC = 0.002 | dAIC = 0.0008 | dAIC = 0.13 | dAIC = 0.03 |
|  |  | w = 10 | - | - | - | - | - |
|  | Mapping quality | MQ = 0 | *ref. level* | *ref. level* | *ref. level* | *ref. level* | *ref. level* |
|  |  | MQ = 30 | **0.28 ± 0.01** | **-0.23 ±0.02** | **-0.21 ±0.02"** | **-0.4 ±0.06** | **-0.13 ±0.02** |

The significance and effect estimates are given for the F1-score (trade-off between precision and recall), the distance between expected and observed community composition (Bray-Curtis and Jaccard distances) and the difference between the observed and expected alpha-diversity estimates. Deviance of the model Akaike Information Criterion (dAIC) is provided for non significant predictors. In other cases, the effect difference between the parameter level and reference level (*ref. level*) is given with standard error (bold stands for *P* < 0.05).

### 2. **Impact of minimap2 mapping options for community structure inference made with the COI barcode**

For the COI barcode, 144 different parameters (three k-mer sizes x three window sizes x four mismatch penalties x two clustering identity thresholds x two mapping quality filters) combinations were considered for each mock community. The impact of these combinations was first evaluated according to the number of false positives and true positive calls. Based on these two criteria, the 97% clustering identity was not considered further as it yielded more false-positive calls and did not increase the true positive calls (supplementary Table 1). Some intermediate values in the other parameters (the mismatch penalty *B* = 3, the window size *w* = 9; supplementary Table 1) were also not considered further as their behaviors were identical to the default values.

Among the remaining 18 sets of parameters considered further (supplementary Table 2), mismatch penalty (*B*) and k-mer window size (*w*) did not affect the F1-score (deviance in model AIC of 0.06 and 0.02 for *B* and *w,* respectively; supplementary Table 3) leaving the k-mer size (*k*) and mapping quality filtering as the most important parameters. The lowest k-mer value of 10 allowed better species identification (increase in F1-score of 0.05 ± 0.02, *P* = 9 x 10^-3^; supplementary Table 3) while increasing mapping quality stringency drastically reduced the F1-score (-0.28 ± 0.02, *P* < 10^-4^).

Mapping quality was also a critical parameter for the community structure determination. Filtering alignment for mapping quality significantly increased the divergence between the observed and expected community structure. This was true using either the Bray-Curtis distance on species relative abundances or the binary presence/absence matrix (*P* < 10^-4^ in both cases), suggesting that the filtered datasets were farther from the true community structure than the unfiltered data (supplementary Table 3). Similarly, filtering alignment for mapping quality reduced alpha diversity (*P* < 10^-4^ for both the Shannon and Simpson index; supplementary Table 3) in agreement with a reduced number of species being detected in this case. The combination of the minimap default values for the mismatch penalty (B = 4) and the window size (w = 10) parameters, with a k-mer size of 10 base pairs and no further filtering on mapping quality (MQ = 0) was deemed as the most optimal pipeline for the COI barcode.
